# Statins Stimulate Hepatic Glucose Production via the miR-183/96/182 Cluster

**DOI:** 10.1101/726695

**Authors:** Tyler J. Marquart, Ryan M. Allen, Mary R. Chen, Gerald W. Dorn, Scot J. Matkovich, Ángel Baldán

**Affiliations:** Edward A. Doisy Department of Biochemistry and Molecular Biology, Saint Louis University, 1100 S. Grand Blvd., Saint Louis, MO 63104; PierianDx Inc., 77 Maryland Plaza, Saint Louis, MO 63108; Department of Medicine, Vanderbilt University Medical Center, 2220 Pierce Ave., 312B PRB, Nashville, TN 37232; Center for Pharmacogenomics, Department of Internal Medicine, Washington University School of Medicine, Saint Louis, MO 63110; Center for Cardiovascular Research, Saint Louis University; Liver Center, Saint Louis University

**Keywords:** SREBP, miRNA, TCF7L2, gluconeogenesis, diabetes

## Abstract

Statins are the most common pharmacologic intervention in hypercholesterolemic patients, and their use is recognized as a key medical advance leading to a 50% decrease in deaths from heart attack or stroke over the past 30 years. The atheroprotective outcomes of statins are largely attributable to the accelerated hepatic clearance of low-density lipoprotein (LDL)-cholesterol from circulation, following the induction of the LDL receptor. However, multiple studies suggest that these drugs exert additional LDL–independent effects. The molecular mechanisms behind these so-called pleiotropic effects of statins, either beneficial or undesired, remain largely unknown. Here we determined the coding transcriptome, miRNome, and RISCome of livers from mice dosed with saline or atorvastatin to define a novel in vivo epitranscriptional regulatory pathway that links statins to hepatic gluconeogenesis, via the SREBP2–miR-183/96/182–TCF7L2 axis. Notably, multiple genome-wide association studies identified *TCF7L2* (transcription factor 7 like 2) as a candidate gene for type 2 diabetes, independent of ethnicity. Conclusion: our data reveal an unexpected link between cholesterol and glucose metabolism, provides a mechanistic explanation to the elevated risk of diabetes recently observed in patients taking statins, and identifies the miR-183/96/182 cluster as an attractive pharmacological candidate to modulate non-canonical effects of statins.

---

Statins are competitive inhibitors of 3-hydroxy-3-methyl-glutaryl-coenzyme A reductase (HMGCR), the rate-limiting enzyme in the cholesterol biosynthetic pathway (1). The ensuing decrease in intracellular sterol levels triggers the proteolytic maturation of the sterol regulatory element-binding protein 2 (SREBP2), which orchestrates a transcriptional program that increases both *de novo* synthesis and uptake of cholesterol. Among the targets of SREBP2, the LDLR is critical in mediating the reduction in plasma LDL-cholesterol which reduces cardiovascular risk in patients taking statins (2). Meta-analyses of epidemiological data and studies in animal models and cells have demonstrated LDL-cholesterol independent effects of statins, both beneficial (improved endothelial function, decreased vascular inflammation, reduced thrombogenic response) and undesired (myalgia, liver damage, memory loss) (3). Importantly, statin therapy is modestly, but significantly, associated dose-dependently with increased new-onset diabetes (reviewed in (4, 5)). Indeed, in 2012 the Federal Drug Administration mandated a modification of the safety label of statins to warn of increased HbA1c and/or fasting plasma glucose. This correlation might be the result of prescription bias, *i.e.* patients who are at increased risk of developing diabetes are also more prone to dyslipidemias and, consequently, are prescribed statins more frequently; however, different laboratories demonstrated decreased pancreatic β-cell insulin secretion (6), peripheral insulin sensitivity (7, 8), and skeletal muscle mitochondrial function (9, 10) following treatment with statins *in vitro* and/or *in vivo*, thus suggesting that statins *per se* do indeed impact glucose homeostasis. The role of statins on hepatic glucose production, however, has never been examined.

MicroRNAs (miRs) are small, noncoding 20-22 nt RNAs that bind to specific, partially complementary sequences in the 3′ untranslated region (UTR) of target mRNAs and recruit them to the RNA-induced silencing complex (RISC) to promote their cleavage and/or translational repression (11). Over the last two decades, miRNAs have been recognized as key regulators of multiple (patho)physiological processes as their expression can be tissue-, developmental-, and disease-specific (12). In the context of cholesterol metabolism, we and others reported on miR-33a (reviewed in (13)): an intragenic miRNA encoded within intron 16 of SREBP2 that fine-tunes the expression of several transmembrane sterol transporters involved in high-density lipoprotein (HDL) lipidation (14–17), bile metabolism (18), and hepatic vesicular secretion (19). Since SREBP2 transactivates its own promoter, miR-33 expression is increased following treatment with statins in cells and *in vivo*. Jeon *et al.* reported that SREBP2 also induces the expression of the miR-183/96/182 cluster and that miR-96 and −182, in turn, target *FBXW7* and *INSIG2* thus providing feed-back regulation for SREBP2 expression and maturation (20). These studies relied on bioinformatic predictions of target genes and exogenous overexpression/inhibition of the microRNA to infer function. However, whether statins actually promote changes in the recruitment of specific mRNAs to the RISC *in vivo* has never been tested.

## EXPERIMENTAL PROCEDURES

### Animal studies

C57BL/6 mice were obtained from NCI, housed in Optimice racks (Animal Care Systems), maintained on a 12h/12h light/dark cycle with unlimited access to food (PicoLab 5353) and water, and bred in the SPF animal facility at Saint Louis University. All mice used for *in vivo* experiments were male at 10–12 weeks of age. Animals used for isolation of primary hepatocytes were male or female. Where indicated, mice were gavaged 100 □L saline or 10 mg/kg atorvastatin (Cayman Chemicals) once a day for ten days, and on the day of sacrifice once at 8 am and fasted for 6 h prior to tissue collection. For overexpression experiments, animals were infused via tail vein with empty or miR-183/96/182 adenoviral vectors (2×10^9^ pfu); pyruvate challenges and tissue collection were done 10- and 12-days post-infusion, respectively. For antisense experiments, animals were gavaged daily with saline or 10 mg/Kg atorvastatin as above for 6 weeks, then kept on drug while injected *i.p.* on days 1, 2, 3, 9, and 12 with 100 □L saline, scrambled, anti-miR-96, anti-miR-182, or anti-miR-183 oligonucleotides (5 mg/kg) or combined anti-miR-96, −182, and −183 oligonucleotides (1.7 mg/Kg each). Pyruvate challenges were done on day 15 at 5 am after removing food at 6 pm the day before. Mice in the antisense experiments were not fasted before collection of blood and livers. For pyruvate challenges, baseline blood glucose was taken from tail snipping using a glucometer (Bayer Contour), then mice were injected *i.p.* with 2 g/kg pyruvate, as described (21), and sequential glucose measurements were taken 15, 30, and 60 min after pyruvate injection. All animals received humane care according to the criteria outlined in the “Guide for the Care and Use of Laboratory Animals” (NIH publication 86-23 revised 1985), and all studies were approved by the IACUC at Saint Louis University.

### MicroRNA vectors

Precursor miRNAs for miR-96, −182, and −183 containing 140 bp upstream and 140 bp downstream of the mature miRNA were amplified from mouse genomic DNA using Platinum Pfx polymerase (Invitrogen), and cloned into pCMV-miR (Origene). To generate replication-deficient adenovirus, a 4,100 bp containing the whole miR cluster (from 140 bp upstream of miR-183 to 140 bp downstream of miR182) was amplified from mouse genomic DNA as above and cloned into pAdTrack-CMV; the vector was then recombined with pAdEasy1, and transfected into Hek293Ad cells to generate high titers of replication-deficient adenovirus, as described (22).

### Anti-miR oligonucleotides

15-mer locked nucleic acid anti-ctrl (5’-TCCTAGAAAGAGTAGA), anti-miR-96 (5’-ATGTGCTAGTGCCAA), anti-miR-182 (5’-TTCTACCATTGCCAA), anti-miR-183 (5’-TTCTACCAGTGCCAT) oligonucleotides were a kind gift from Miragen Therapeutics Inc. Antimirs were resuspended in sterile saline and stored at −20°C until used.

### Primary hepatocytes

Cells were isolated from 8–10-week-old, male C57BL/6 mice fed chow, using Perfusion and Digest buffers (Invitrogen). Cells were resuspended in William’s E Medium (Invitrogen) supplemented with Plating Supplements (Invitrogen), plated in 12- or 6-well BioCoat Collagen I plates (BD), and incubated at 37°C and 5% CO_2_ for 6 h. Then, the media was switched to William’s E supplemented with Maintenance Supplements (Invitrogen). Where indicated, cells were cultured in media supplemented with 1 μmol/L atorvastatin or pravastatin, 5 μmol/L simvastatin (Cayman), or 10 nmol/L siRNA-control or siRNA-Tcf7l2 (Dharmacon D-001810 and L-058892, respectively). Glucose production assays were performed as described (23). Briefly, cells were incubated 6 h in 1 mL glucose production buffer (glucose-free DMEM without phenol red supplemented with 20 mmol/L sodium lactate and 2 mmol/L sodium pyruvate. Supernatants were collected and used to measure glucose with a colorimetric kit (Wako), and data were normalized to intracellular protein.

### Luciferase reporter assays

HEK293 cells were transiently transfected with luciferase reporters and miR expression plasmids in triplicate in 24-well plates using the calcium phosphate method. Cell extracts were prepared 48 h later, and luciferase activity measured using the Luciferase Assay System (Promega). Data were normalized to β-galactosidase activity to correct for small changes in transfection efficiency.

### RNA analysis

RNA was extracted from mouse livers and cells using Trizol (Invitrogen). RISC-associated RNA was immunoprecipitated from liver homogenates using an anti-AGO2 antibody (Wako, clone 2D4) and extracted as described (24). Bar-coded libraries for small and polyA-enriched large RNAs, and for RISC-associated RNAs were prepared for livers from saline (n=8) and atorvastatin (n=8) treated mice using Illumina kits, and transcriptional profiling by deep sequencing were analyzed as described (24, 25). For coding transcriptome, data from Saline-4 and Atorvastatin-2 libraries were discarded due to poor sequencing. For miRNome, all 16 libraries produced good quality data. For RISCome, libraries for Saline-1, −3, −4, −5, and 6, and for Atorvastatin-3, −4, −5, and −7 produced quality data. Annotated sequencing data are available in Supplemental Files 1–3. Predicted miRNA–mRNAs pairs/networks were evaluated with miRTarVis, as described (26). For all other experiments, cDNAs were generated from 1 μg of DNase1-treated RNA using Superscript III (Invitrogen) and random hexamers. Quantitative real-time PCR was done with Power SybrGreen reagent (Applied Biosystems), using a LightCycler-480 (Roche). Primer sets are available upon request. Values were calculated using the comparative ΔΔC_t_ method and normalized to ribosomal *Rplp0*. Targeted miRNA expression was analyzed using miRCURY LNA reagents and assays (Exiqon) and normalized to U6.

### Protein analysis

Protein were extracted from cells or livers using HEPES buffer (20 mmol/L HEPES pH 7.4, 150 mmol/L NaCl, 0.2 mmol/L EDTA, 2 mmol/L MgCl_2_, 1% Triton X-100, 10% glycerol) containing protease inhibitor cocktail (Pierce). Forty micrograms of protein were resolved in 4–12% polyacrylamide gels, transferred to PVDF membranes, and probed with antibodies for TCF7L2 (1:1,000 dilution, Cell Signaling C48H11), PCK1 (1:1,000 dilution; Abcam ab70358), G6PC (1:1,000 dilution; Novus NBP1-80533) and β-ACTIN (1:5,000 dilution; SCBT sc-130656) in TBS-Tween20 containing 4% non-fat dry milk. Immune complexes were detected with horseradish peroxidase-conjugated anti-rabbit secondary antibodies (1:5,000 dilution; Bio-Rad 1706515).

## RESULTS

In an effort to understand the functional changes in the hepatic transcriptome that follow treatment with statins, we dosed C57BL/6 mice with saline or atorvastatin, and use RNA-seq to define total mRNAs (coding transcriptome), mRNAs associated with the RISC (RISCome), and miRNAs (miRNome) (Fig. 1a). The complete annotated sequencing data are available in Supplementary Files 1–3. Fig. 1b shows that ~10% of the coding transcriptome changed in response to the drug. As expected, at least 15 canonical targets of SREBP2 such as *Hmgcr,* squalene synthase (*Fdft1*), lanosterol synthase (*Lss*), 7- and 24-dehydrocholesterol reductases (*Dhcr7*, *Dhcr24*), proprotein convertase subtilisin/kexin type 9 *(Pcsk9)*, and LDL-receptor (*Ldlr*) were induced in the livers of atorvastatin-treated mice, compared to saline (Supplementary File 1). Data in Figs. 1c and 1d, and in Supplementary Files 2 and 3 show that 62 miRNAs and 1,504 RISC-associated mRNAs were differentially regulated by the statin. Our ultimate goal, however, was to identify transcripts that were both reciprocally regulated in total transcriptome vs. RISCome, and predicted as targets of miRNAs differentially regulated in the miRNome. Analysis of the three sets of data revealed several miR/mRNA pairs that fit those criteria (Fig. 1e and Supplemental Data Fig. 1). In the rest of this report, we focus on the miR-183/96/182 polycistronic cluster, because all 3 miRNAs were upregulated in the livers of mice treated with atorvastatin, compared to saline (Fig. 1c), and 9 predicted targets for one or more miRNAs in that cluster were both RISC-enriched and transcriptome-depleted in the same livers (Fig. 1e). These targets are *Bicd2* (bicaudal D homolog 2), *Mkl2* (myocardin-like 2), *Nox4* (NADPH oxidase 4), *Pik3r1* (phosphoinositide-3-kinase regulatory subunit 1), *Pxmp4* (peroxisomal membrane protein 4), *Sfxn5* (sideroflexin 5), *Slc22a23* (solute carrier family 22 member 23), *Tcf7l2* (transcription factor 7 like 2), and *Tmem189* (transmembrane protein 189). Data in Figs. 2a and 2b show the relative abundance of miR-183, −96, and −182 in the livers of saline-treated mice, and the fold-induction following treatment with saline. We also highlight miR-33 in the same panels, as we and others have previously described this miRNA as regulated by statins. Data in Fig. 2c show the transcriptome and RISCome scores for the nine potential targets defined above. Importantly, both the induction of the miR cluster and the repression of the nine putative targets in response to statins were validated by RT-qPCR in a second cohort of mice (data not shown) and in mouse primary hepatocytes (Fig. 2d). Collectively, data in Figs. 1 and 2 demonstrate that miR-183/96/182 is an *in vivo* hepatic sensor of exposure to statins, and that these miRNAs actively recruit specific mRNA targets to the RISC in response to statins.

**Figure 1.**
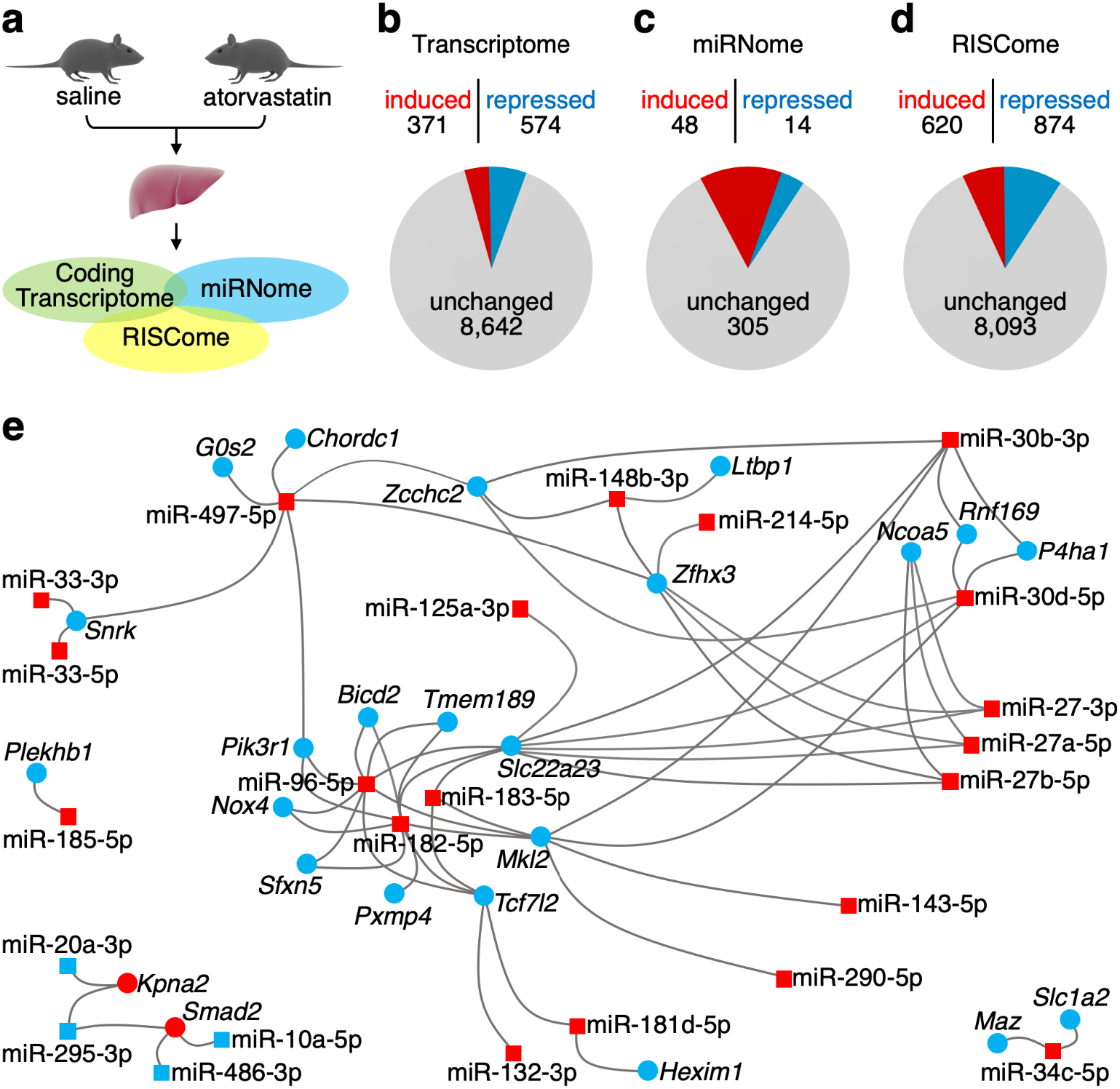
Atorvastatin-dependent changes in hepatic transcriptome, miRNome, and RISCome. **a,** Experimental design. Mice were gavaged daily with saline or 10 mg/Kg atorvastatin. See Methods for details on RNA analysis. **b,** Summary of changes in total coding transcriptome in atorvastatin (*n*=7) vs. saline (*n*=7). **c,** Summary of changes in miRNome (*n*=8/group). **d,** Summary of changes in RISCome (*n*=5-saline; n=4-ATR). **e,** Node-link diagram of miRTarVis-predicted miRNA–mRNA networks changing in atorvastatin vs. saline; upregulated transcripts in red, downregulated transcripts in blue. See Supplemental Files 1–3 for complete annotated sequencing data and analysis.

**Figure 2.**
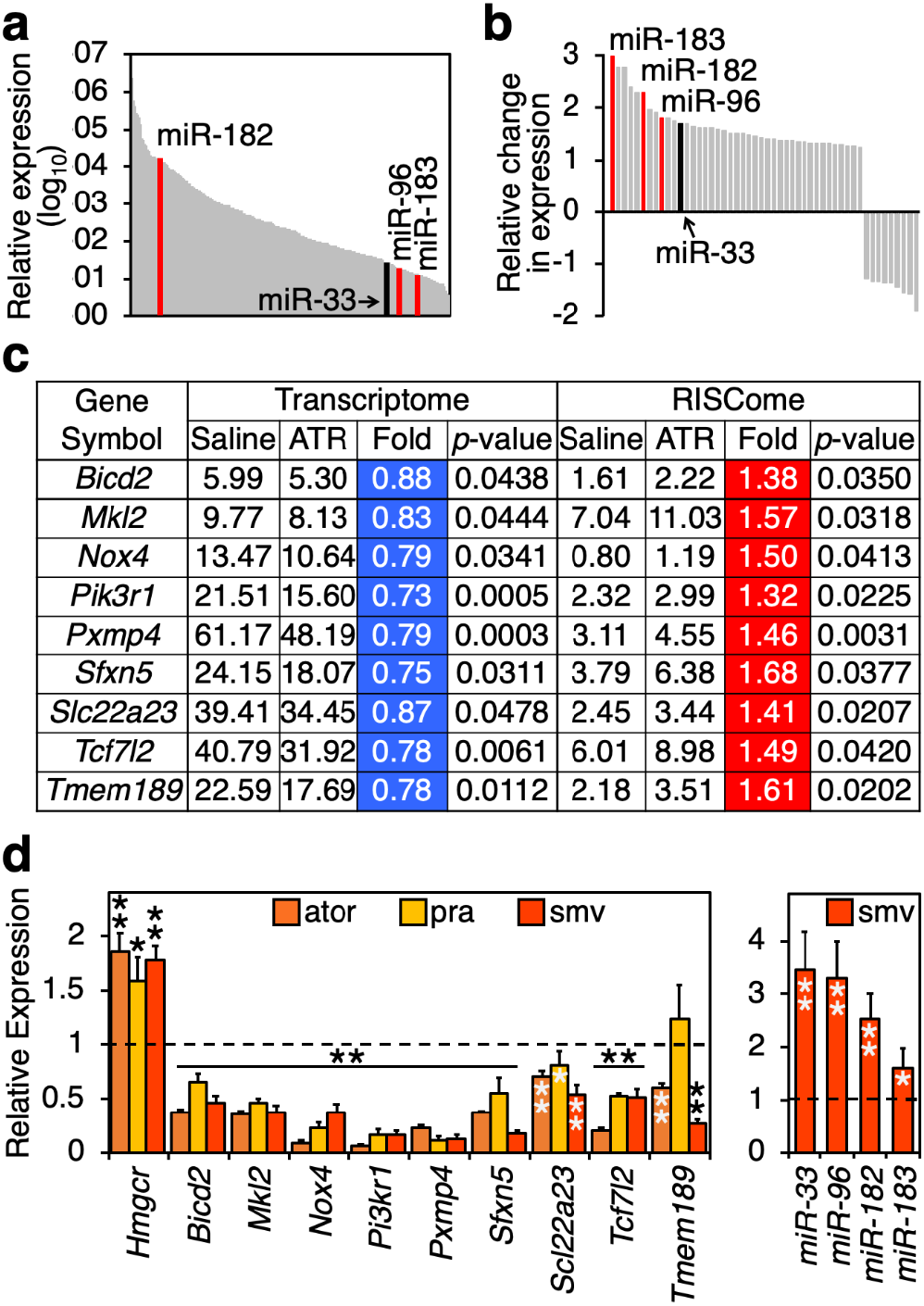
Treatment with statins induces the recruitment of miR-183/96/182 targets to the RISC. **a,** Relative abundance of mature miR-183, −96, and −182 in the hepatic miRNome of saline-treated mice. For comparison, the previously reported statin-regulated miR-33 is shown in black. **b,** Relative fold-induction of mature miR-183, −96, and −182 in response to atorvastatin. **c,** Nine putative miR-183/96/182 targets identified in Fig. 1e were simultaneously depleted in total transcriptome and enriched in the RISCome of mice dosed with atorvastatin. **d,** Validation by qPCR of changes in hepatic miR-183/96/182 and their putative targets in a second cohort of mice (n=5) dosed with saline or atorvastatin, as in Figs. 1. and 2a–c. Canonical SREBP2 targets (*Hmgcr*, *Ldlr*, *Pcsk9,* and *miR-33* are shown as positive controls). **e,** Validation of changes in miR-183/96/182 and their putative targets in mouse primary hepatocytes cultured 24 h in media supplemented with 1 μmol/L atorvastatin (ator) or pravastatin (pra), or 5 μmol/L simvastatin (smv) (n=3/group). Canonical SREBP2 targets *Hmgcr* and *miR-33* are shown as positive controls. Dotted line represents normalized expression in dmso-treated cells. \**p*<0.05, \*\**p*<0.01, compared to dmso.

In agreement with a report by Jeon *et al.* (20), we mapped two SREBP-responsive elements (SRE) upstream of the transcription start for the miR-183/96/182 cluster (Supplemental Data Fig. 2a). The proximal element is evolutionarily conserved between mice and humans, while the distal one is divergent (Supplemental Data Fig. 2b). Nevertheless, both sequences in the human and mouse promoters confer responsiveness to SREBP2 (Supplemental Data Fig. 2c, d).

Of the nine transcripts that originally fit our criteria (Figs. 1e and 2c), we decided to focus on *Tcf7l2*, as it has been identified as a key transcriptional regulator of hepatic gluconeogenesis, and polymorphisms in this locus have been associated with diabetes in several GWAS (27–31). We hypothesized that statins stimulate hyperglycemia, at least in some patients, by increasing hepatic glucose output via miR-183/96/182–TCF7L2. The miRNA target prediction algorithm Targetscan (32) identifies conserved sequences in the 3’ UTR of *TCF7L2* that are partially complementary to miR-183, −96, -or 182 (Fig. 3a). These sequences, or the entire 3’ UTR of mouse or human *TCF7L2*, were cloned downstream of a luciferase reporter and transiently transfected into HEK293 cells in the presence or absence of a plasmid encoding each miR. Data from these experiments in Fig. 3b show that each isolated conserved sequence (but not sequence 1 in human *TCF7L2*, which is not conserved in mouse), as well as the whole 3’ UTR, were able to confer responsiveness to miR-183, −96, and −182. Additionally, point mutations in each predicted sequence to prevent interaction with the seed sequence of each miRNA abrogated the response (Supplemental Data Fig. 3). Importantly, data in Fig. 3c using anti-miR oligonucleotides demonstrate that in mouse primary hepatocytes the miR-183/96/182 cluster mediates the repressive effect of atorvastatin on *Tcf7l2.* Together, data in Fig. 3 identify conserved sequences in *TCF7L2* as functional statin and miR-183/96/182 responsive elements.

**Figure 3.**
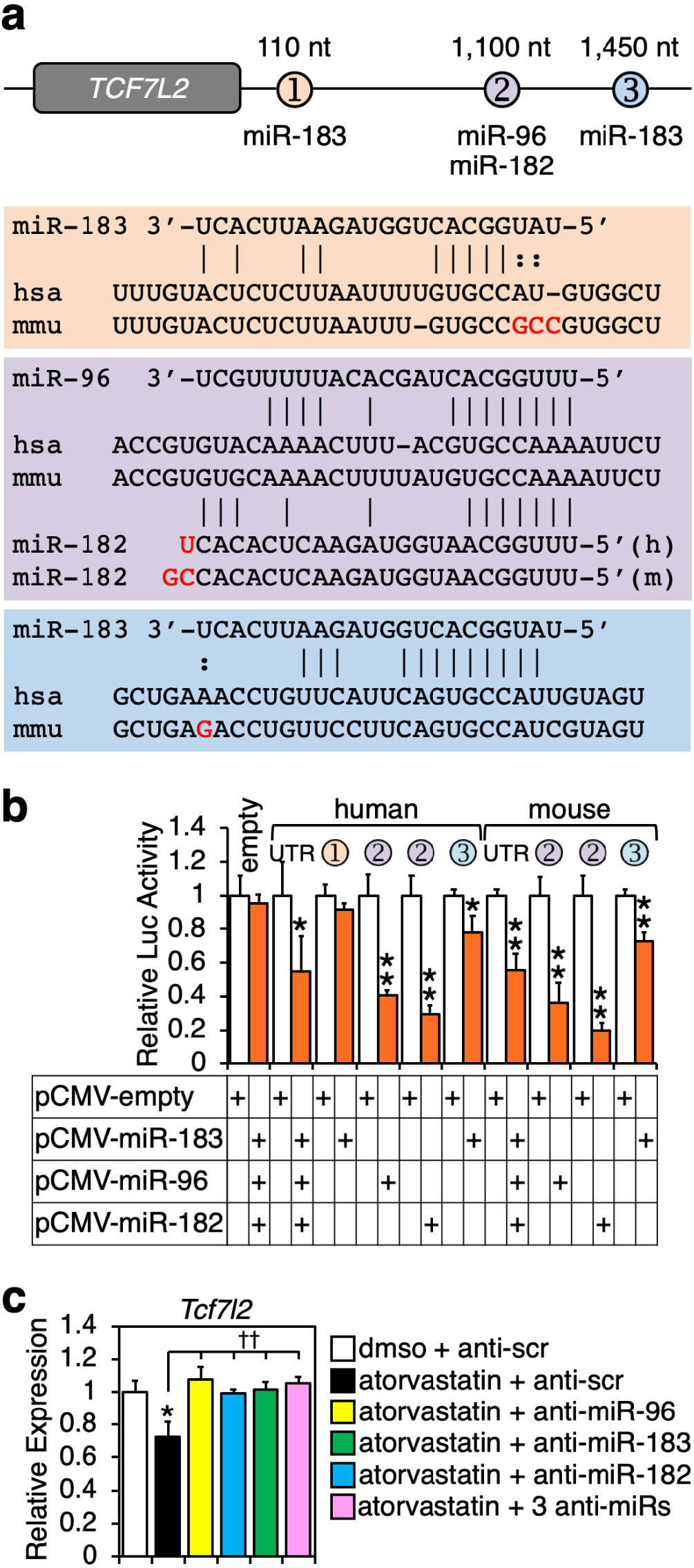
Functional miR-183/96/182-reponsive elements in *TCF7L2*. **a,** Conserved sequences in the 3’UTR of *TCF7L2* with partial complementarity to miR-183/96/182 are color-coded as elements 1, 2, and 3. Distance from the stop codon are indicated. Alignments for mouse and human sequences are shown. Solid lines represent Watson-Crick complementarity; colon marks represent divergence between mouse and human sequences. **b,** Normalized luciferase activity in extracts from HEK293 cells transiently transfected with a reporter fused to the potential miR-responsive elements identified above or the whole 3’UTR, and expression vectors for each miR. As negative control, we used the empty pGL3Promoter vector. Data are mean ± SE of 3 independent experiments in quadruplicate. \*\**p*<0.01. **c,** miR-183/96/182 mediate atorvastatin-induced repression of *Tcf7l2*. Mouse primary hepatocytes were transfected with scrambled or anti-miR oligonucleotides, and 24 h later switched to media supplemented with vehicle or atorvastatin for 24 h. Data are mean ± SE of 2 independent experiments in quadruplicate. \**p*<0.05, compared to dmso; ^††^*p*<0.01, compared to atorvastatin plus anti-scr.

We next tested the ability of statins, the miR cluster, and TCF7L2 to modulate glucose production in mouse primary hepatocytes. Data in Fig. 4 show that gluconeogenesis was increased in cells incubated with either atorvastatin, simvastatin, or pravastatin, compared to vehicle (Fig. 4a); in cells overexpressing miR-183/96/182 (via adenoviral transduction), compared to empty vector (Fig. 4b); and in cells where *Tcf7l2* was silenced using siRNA, compared to control oligonucleotides (Fig. 4c). Consistent with these data from cultured hepatocytes, the amounts of the key gluconeogenic enzymes phosphoenolpyruvate kinase 1 (PCK1) and glucose-6-phosphatase, catalytic subunit (G6PC) increased in the livers dosed with atorvastatin, compared to saline, and were inversely correlated to those of TCF7L2 (Fig. 4d). Collectively, these data demonstrate that statins, the miR cluster, and TCF7L2 are critical regulators of the gluconeogenic pathway in cultured hepatocytes.

**Figure 4.**
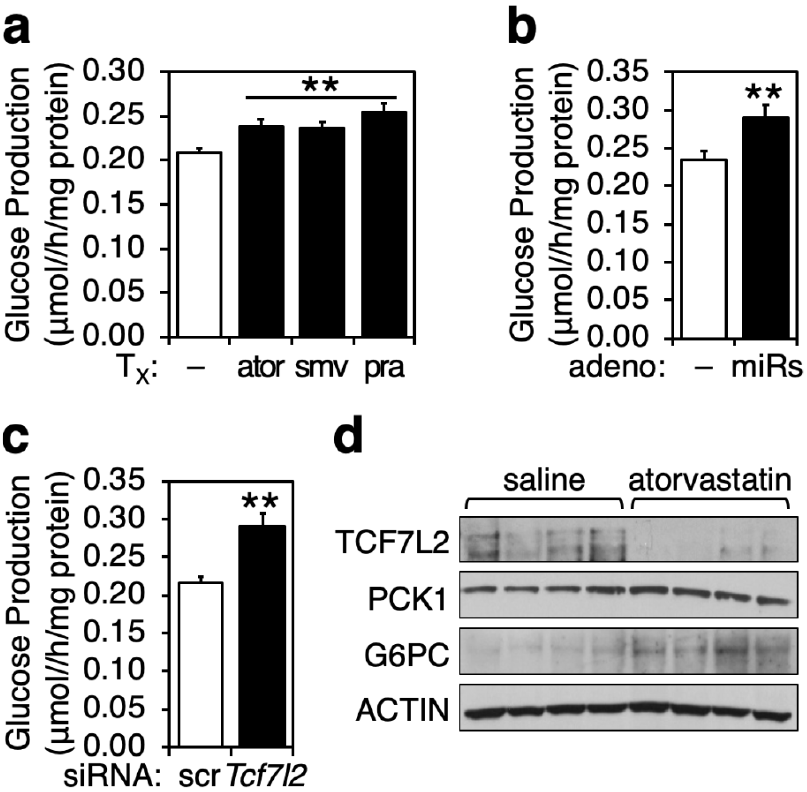
Statins, miR-183/96/182, and TCF7L2 modulate hepatic gluconeogenesis. **a,** Glucose production in mouse primary hepatocytes incubated in media supplemented with vehicle, 1 μmol/L atorvastatin, 5 μmol/L simvastatin, or 1 μmol/L pravastatin. **b,** Glucose production in mouse primary hepatocytes after transduction with empty adenovirus or adenovirus encoding the miR-183/96/182 cluster. **c,** Glucose production in mouse primary hepatocytes after transfection with siRNA-control or siRNA-Tcf7l2 oligonucleotides. **d,** Hepatic protein abundance of TCF7L2 and gluconeogenic enzymes PCK1 and G6PC in mice dosed with saline or 10 mg/Kg/d atorvastatin for 6 weeks. Data in panels a–c are mean ± SE of at least 2 independent experiments in quadruplicate. \*\**p*<0.01

To test the functional consequences of miR-183/96/182 overexpression on hepatic glucose production *in vivo*, we transduced mice with empty adenovirus or adenovirus encoding the miR cluster. After 10 days, animals were fasted overnight and then injected *i.p.* with pyruvate early in the morning. Hepatic gluconeogenic capacity was estimated from the conversion of pyruvate to glucose and its release into circulation, which was monitored with a standard glucometer. Data in Fig. 5a and Supplemental Data Fig. 4 show that, in two independent experiments, mice overexpressing the miR cluster were able to sustain increased glycemia in response to the pyruvate challenge, compared to control animals. These data demonstrate increased gluconeogenic capacity in the former mice. Animals were allowed to recover after the pyruvate test for 48 h, and then some were fasted overnight, and some allowed access to food. In agreement with the pyruvate challenge results, both mRNA (Fig. 5b and Supplemental Data Fig. 4b) and protein (Fig. 5c) panels show that miR overexpression in the liver resulted in the decrease of TCF7L2 and the concomitant increase in PCK1 and G6PC. These results are consistent with the previously reported suppressive effect of TCF7L2 on the gluconeogenic pathway via negative regulation of *PCK1* and *G6PC* expression (see discussion below). Interestingly, these changes were much more apparent in the livers of fed mice, and indeed PCK1 and G6PC proteins were as robustly expressed (if not more) in the livers of fed miR-overexpressing mice as in fasted control livers (Fig. 5c). This latter observation suggests that the miR-183/96/182–TCF7L2 pathway can overcome the canonical nutritional regulation of these gluconeogenic genes.

**Figure 5.**
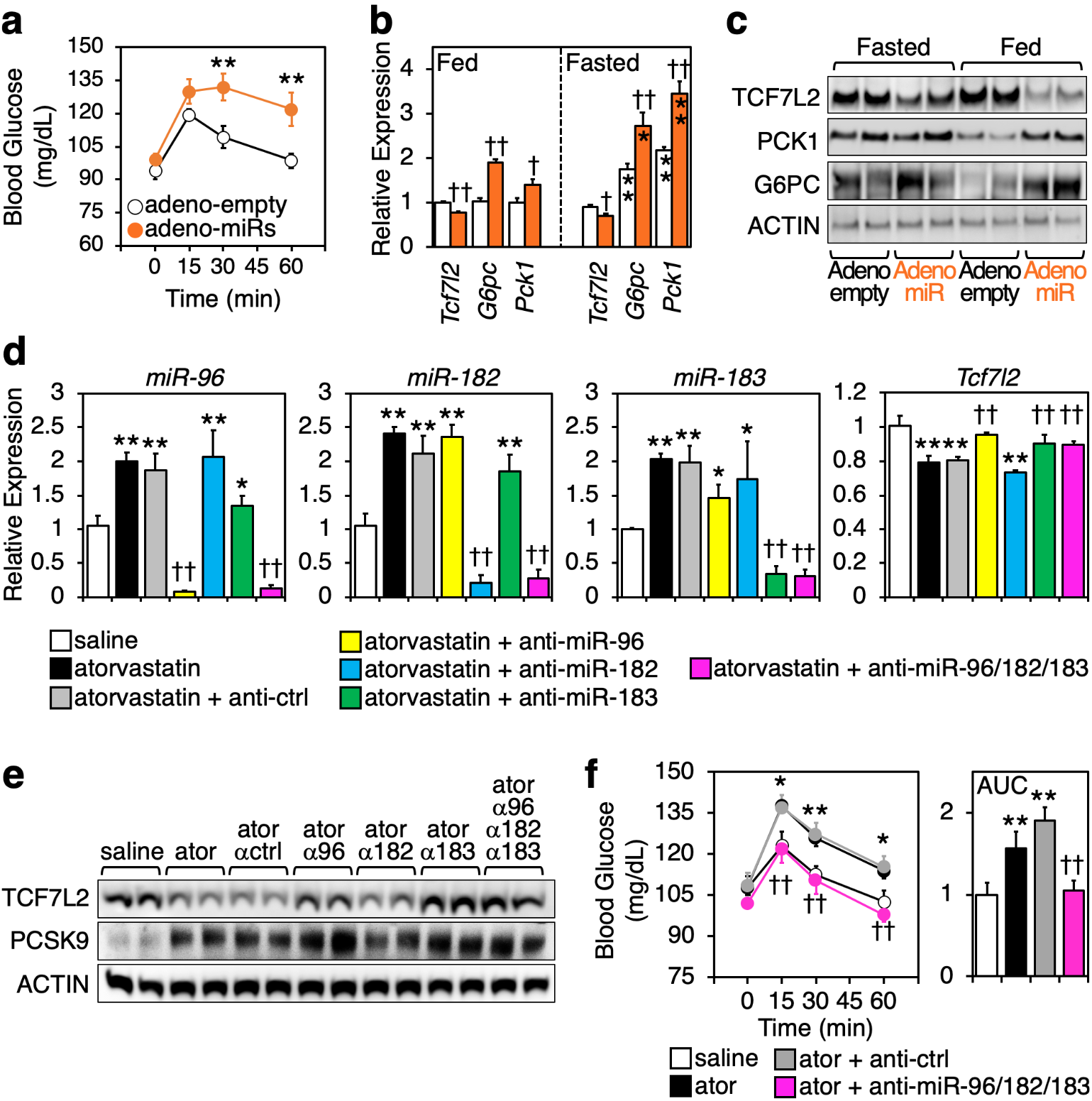
Therapeutic silencing of the miR-183/96/182 cluster abrogates statin-induced hepatic gluconeogenesis. **a,** Blood glucose curves in response to pyruvate challenge (see Methods) show increased hepatic glucose output in mice transduced with adenovirus encoding the miR cluster, compared to empty adenovirus. Data are mean ± SE; *n*=10/group; \*\**p*<0.01. **b,** Hepatic protein levels for TCF7L2, PCK1, and G6PC two days after the pyruvate challenge, in mice allowed access to food, or fasted overnight. Data are mean ± SE; n=5/group; \**p*<0.05, \*\**p*<0.01, compared to fed mice; ^†^*p*<0.05, ^††^*p*<0.01, compared to empty adenovirus. **c,** Immunoblots of hepatic protein extracts in the same mice as (b). **d,** Mice were gavaged with saline or 10 mg/Kg/d atorvastatin daily for 6 weeks, then treated with control or anti-miR oligonucleotides, alone or combined, for 15 days (see Methods). Atorvastatin induces the hepatic expression of the 3 miRs and reduces that of *Tcf7l2*. Anti-miRs are specific in silencing the expression of their cognate miR. Silencing miR-96 or −183 abolished atorvastatin-induced repression of *Tcf7l2*. Data are mean ± SE; *n*=6/group; \**p*<0.05, \*\**p*<0.01, compared to saline; ^†^*p*<0.05, ^††^*p*<0.01 compared to atorvastatin plus anti-ctrl. **e,** Changes in TCF7L2 protein levels parallel those in mRNA. PCSK9 was used as positive control to monitor the efficacy of the statin. **f,** Hepatic gluconeogenic capacity in response to a pyruvate challenge was tested in the 4 groups shown, confirming the functional role of the miR cluster as the mediator to statin-induced hepatic glucose production. Area under the curve represented in arbitrary units. Data are mean ± SE; *n*=6/group; \**p*<0.05, \*\**p*<0.01, compared to saline; ^††^*p*<0.01, compared to atorvastatin plus anti-ctrl.

Finally, we tested whether therapeutic silencing of each miR (or of all three) could revert the atorvastatin-induced increase in hepatic gluconeogenesis. Mice were randomized for daily gavage with saline or atorvastatin during 6 weeks, and then kept on treatment while dosed with one or more anti-miR oligonucleotides on days 1, 2, 3, 9 and 12 (see Methods for details). Data in Fig. 5d show that the statin (or statin plus scrambled anti-miR), raised the expression of all three miRNAs in liver, and that the anti-miRs were specific in silencing the expression of each cognate miR. Interestingly, silencing miR-96 modestly reduced the expression of miR-183, and vice-versa. This is consistent with reports in other multi-cistronic miRNAs where altering the expression of one miR affects the levels of the others in the cluster. Nonetheless, *Tcf7l2* mRNA (Fig. 5d) and protein (Fig. 5e) levels were efficiently reduced in the livers of mice dosed with the statin, or statin plus scrambled anti-miR. Importantly, silencing miR-96 or miR-183, but not miR-182, negated the statin-induced reduction in *Tcf7l2* both at the mRNA (Fig. 5d) and protein (Fig. 5e) level. Silencing all three miRNAs did not provide further increase in TCF7L2 abundance. Supplemental Data Fig. 5a show that the expression of *G6pc* in the same livers largely opposed that of *Tcf7l2*: both gluconeogenic transcripts were induced by the statin, and treatment with anti-miRs abrogated the statin-dependent induction. Together, these data demonstrate that the miR-183/96/182 cluster is the *in vivo* mediator of the statin-induced regulation of hepatic TCF7L2 and gluconeogenic enzymes. Interestingly, anti-miR182 was able to reduce the expression of *G6pc* (Supplemental Data Fig. 5a), despite having no effect on *Tcf7l2* levels (Fig. 5c, d). No miR-182 binding sites are predicted in murine or human *G6PC*. Thus, the mechanism by which miR-182 modulates *G6pc* expression independent of *Tcf7l2* remains unknown. Supplemental Data Fig. 5a also shows results for the other 8 transcripts initially identified in this study: data demonstrate that the miR-183/96/182 cluster does indeed control the liver expression of *Sfxn5*, *Tmem189*, *Slc22a23*, *Nox4*, and *Mkl2*; however, no changes in the mRNA amounts of *Pi3kr1*, *Pxmp4*, and *Bicd2* were noted. The reasons for the lack of response to atorvastatin of these latter genes are unclear, since both transcripts clearly were regulated in other *in vivo* (Fig. 1) and in culture (Fig. 2d) experiments. It is possible that the nutritional status of the mice (animals were not fasted in Fig. 5d–f and Supplemental Data Fig. 5, as opposed to those in Figs. 1 and 2) may contribute additional regulatory mechanisms, beyond SREBP2, for these genes. Nonetheless, Fig. 5f shows data from the same animals two days before sacrifice, when they were challenged with pyruvate to measure *in vivo* gluconeogenesis. Consistent with the data in Fig. 5a from miR-overexpressing mice, hepatic glucose production was induced in mice dosed with atorvastatin (or atorvastatin plus scrambled anti-miR), compared to animals receiving saline; importantly, treatment with anti-miRs abrogated the effect of the statin and returned glycemic kinetics to control levels. These latter results upon pyruvate challenge were reproducible in a second cohort of mice treated as above (Supplemental Data Fig. 5b). Collectively, data in Fig. 5 demonstrate that statins promote hepatic gluconeogenesis via miR-183/96/182.

## DISCUSSION

The cardiovascular therapeutic benefits of statins are presumed to outweigh their diabetogenic effect. Yet, it is critically important to establish the molecular mechanisms behind this undesired side effect. This is especially important when considering preventive prescription to younger dyslipidemic patients who will presumably be taking statins for several decades. Clinical and *in vitro* studies showed that statins impair both pancreatic insulin secretion and peripheral insulin sensitivity (4, 5). Ours is the first report demonstrating a direct effect of statins on hepatic glucose production. We show that the SREBP2–miR-183/96/182–TCF7L2 axis controls the expression of gluconeogenic enzymes, and increases glucose production both in cultured hepatocytes and in mice (Fig. 6). Additionally, we show that therapeutic silencing of the miR cluster normalizes *Tcf7l2* expression and reverts statin-stimulated hepatic glucose production. TCF7L2 is a member of the Wnt signaling pathway (33), and modulates the transcription of target genes by interacting not only with β-catenin (its canonical partner), but also with the key metabolic transcription factors FOXO1 (34) and HNF4α (35). The exact role of TCF7L2 on glucose metabolism has been controversial for some time (36), and authors suggested that TCF7L2 acts as a transcriptional activator or repressor depending on its partner. In addition, at least 13 different transcripts and 2 different sized proteins are expressed from *TCF7L2*, and it is unclear whether these isoforms differ in their trans-activating/repressing properties. Four different *Tcf7l2*^−/−^ models have been generated, all with perinatal lethality(34, 37–39). Oh *et al.* showed that *Tcf7l2*^+/−^ mice display decreased glucose tolerance and insulin sensitivity, and exhibited increased hepatic expression of gluconeogenic genes (34). In contrast, Savic *et al.*(38) and Boj *et al.*(39) reported improved glycemic control using two other heterozygote mouse models and a BAC transgenic mouse, but did not report on hepatic glucose production. The reasons for the discrepancies among these different genetic models remain unknown. More recently, a liver-specific transgenic mouse expressing a dominant negative form of TCF7L2 showed progressive impairment in response to pyruvate challenge in the absence of hepatic insulin intolerance (40). Collectively, these studies show that TCF7L2 activity modulates hepatic and peripheral glucose metabolism, and whole-body glycemic control. In regards to gluconeogenesis, the literature strongly suggests that TCF7L2 reduces hepatic gluconeogenesis likely by decreasing the transcriptional activity of positive regulators of *PCK1* and *G6PC (34*–*36*, *41*, *42)*.

**Figure 6.**
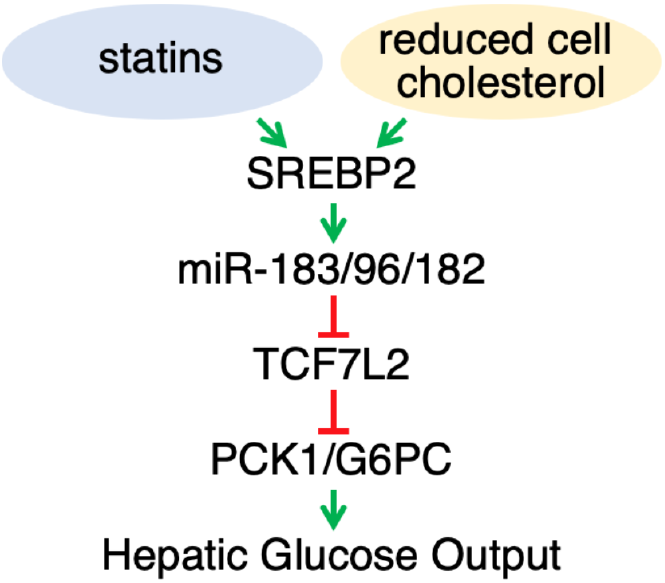
The SREBP-2–miR-183/96/182–TCF7L2 modulates hepatic glucose output. During episodes of low-intracellular cholesterol, such as in response to statins, the expression of miR-183/96/182 is induced, which in turns reduces the levels of gluconeogenic repressor TCF7L2, ultimately promoting accelerated hepatic glucose production.

Interestingly, *TCF7L2* transcription is stimulated postprandially by insulin (41). One key observation in our studies is that the induction miR-183/96/182 overrides the canonical nutritional control of the gluconeogenic program (on during fasting, off postprandially). As a consequence, we demonstrate that both the mRNA and protein amounts of both PCK1 and G6PC are similar in fed miR-overexpressing livers than in fasted control livers. This has particularly worrisome implications when considering patients taking statins for extended periods of time, whom presumably will have persistently elevated expression of the miR cluster: our results suggest that these patients may have sustained activation of the gluconeogenic pathway. In an otherwise healthy individual, this accelerated glucose delivery from liver into circulation should trigger homeostatic insulin-dependent clearance in peripheral tissues to maintain euglycemia. However, it is tempting to speculate that the sustained unregulated hepatic glucose production in response to statins may over time elevate blood HbA1c and fasting glucose, and ultimately contribute to new-onset diabetes in “vulnerable” patients. Meta-analyses of clinical trials and population-based studies revealed statin therapy increases the risk of new-onset diabetes up to 43%. However, why only some, but not all, patients taking statins develop diabetes remains a mystery. Additional research is necessary to determine whether these “vulnerable” patients, who are prone to develop statin-induced diabetes, have polymorphisms in the *miR-183/96/182* or *TCF7L2* loci that make them more sensitive (*i.e.*, hyper-responsive) to statins.

## ACKNOWLEDGMENTS

We thank Dr. Aimee Jackson at miRagen Therapeutics Inc. for providing the anti-miR oligonucleotides for *in vivo* experiments. We also thank Dr. Cedric Langhi at NovoNordisk for discussions. This study was supported in part by NIH HL107794 and AHA GRNT20460189 grants (to Á.B.), Washington University Department of Medicine start-up funds (S.J.M.), and AHA PRE7240026 fellowship (to R.M.A.).

## AUTHOR CONTRIBUTIONS

T.J.M. and Á.B. conceived the idea, designed experiments, and wrote the manuscript. T.J.M. and R.M.A. performed experiments, and collected and analyzed data. M.R.C. analyzed data. G.W.D. and S.J.M. supervised library construction, sequencing, and bioinformatics analysis.

