## Supplemental figures for "Statins Stimulate Hepatic Glucose Production via the miR-183/96/182 Cluster"

**SUPPLEMENTAL FIGURES 1–5**

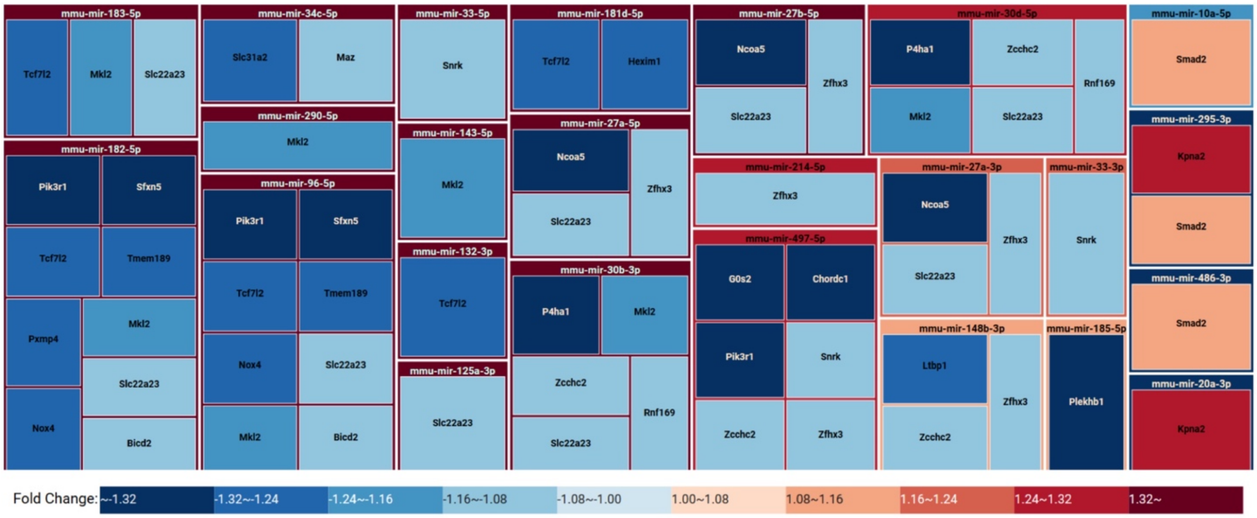


**Supplemental Figure 1. miRTarVis predicted miRNA-mRNA pairs regulated by atorvastatin.** Data are the same as in node-link diagram in Fig. 1e, but here are shown as a heatmap table.

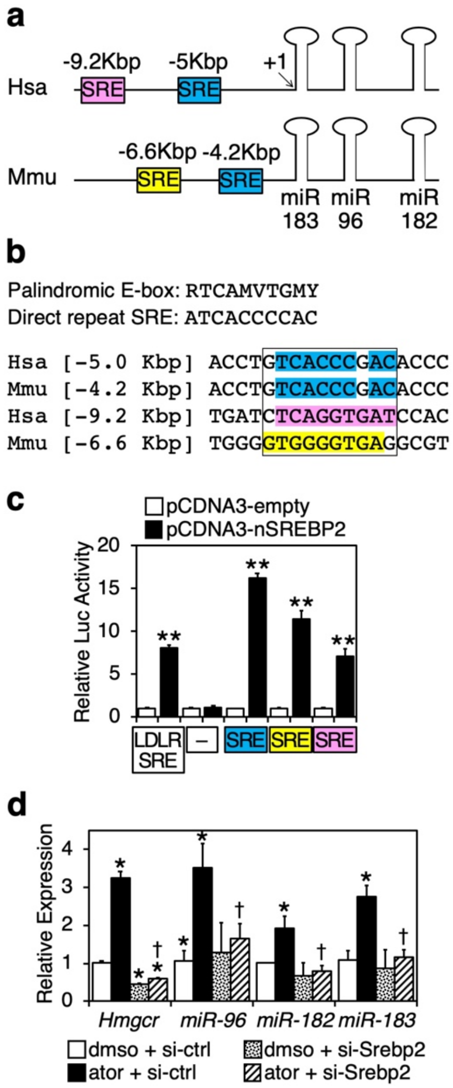


**Supplemental Figure 2. SREBP2 mediates statin-induced expression of the miR-183/96/182 cluster. a,** Putative SREBP responsive sequences upstream of the human and murine miR cluster. +1 denotes the first nucleotide in mature miR-182. **b,** The proximal SRE sequences are identical between human and mouse, while the distal SRE are divergent. **c,** Reporter luciferase assays in Hek293 cells transiently transfected with reporter plasmids containing the different SREs in the mouse and human miR-183/96/182 promoters. An empty luciferase reporter and a reporter containing the SRE from human LDLR were used as a negative and positive controls, respectively. **d,** Mouse primary hepatocytes were transfected with siRNA-ctrl or siRNA against *Srebp2*, and 24 h later cultured in media supplemented with or without 1 μmol/L atorvastatin. The relative expression of the miRs was analyzed by qPCR 24 h later. **p*<0.05, compared to DMSO + siRNA-ctrl. †*p*<0.05, si-Srebp2 compared to si-ctrl.

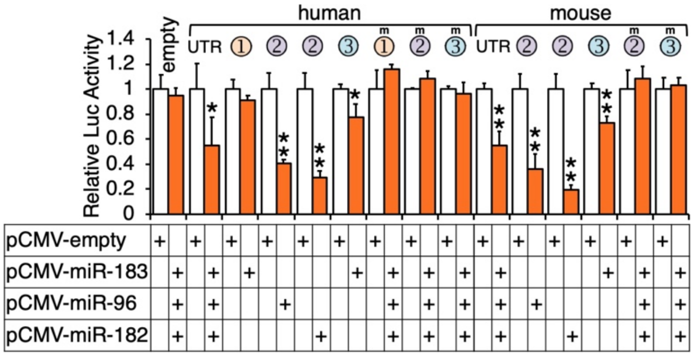


**Supplemental Figure 3. Conserved sequences in the 3'UTR of *TCF7L2* confer response to miR-183/96/182.** Expanded data for Fig. 2b containing point mutants (m) for each of the putative response sequences in human and mouse *TCF7L2*. Normalized luciferase activity in extracts from Hek293 cells transiently transfected with the indicated reporters and miR expression plasmids. **p*<0.05, ***p*<0.01 compared to pCMV-empty.

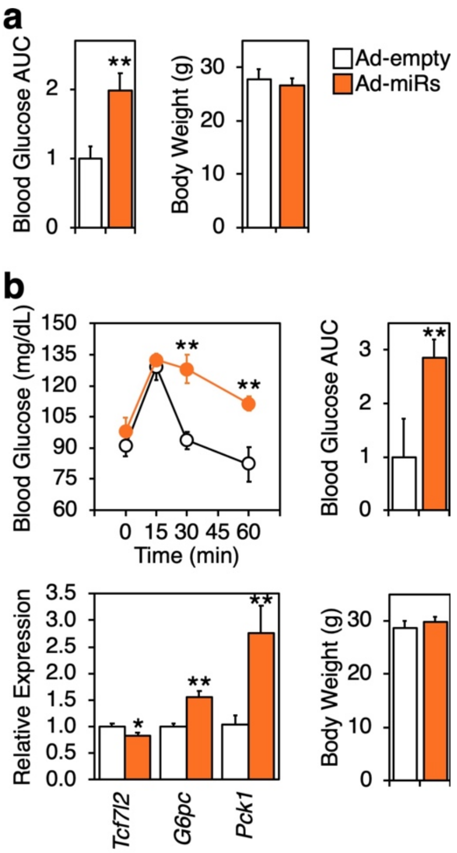


**Supplemental Figure 4. Hepatic overexpression of miR-183/96/182 promotes gluconeogenesis. a,** Area under the curve (AUC) for the pyruvate challenge experiment shown in Fig. 4a. **b,** A second cohort of mice was transduced via tail vein injection with empty adenovirus or adenovirus encoding miR-183/96/182 (n=6/group). Pyruvate challenge was performed 10 days post-infusion, as in Fig. 4a. Increased gluconeogenic capacity was evident in mice overexpressing the miR cluster, as shown by increased blood glucose and AUC, and elevated expression of hepatic *G6pc* and *Pck1*. Consistent with previous figures, hepatic *Tcf7l2* expression was reduced in the same mice overexpressing the miR cluster. No changes in body weight were noted in either the first or the second cohort of mice. Data are mean ± SE. *$p$<0.05, **$p$<0.01, compared to empty adenovirus.

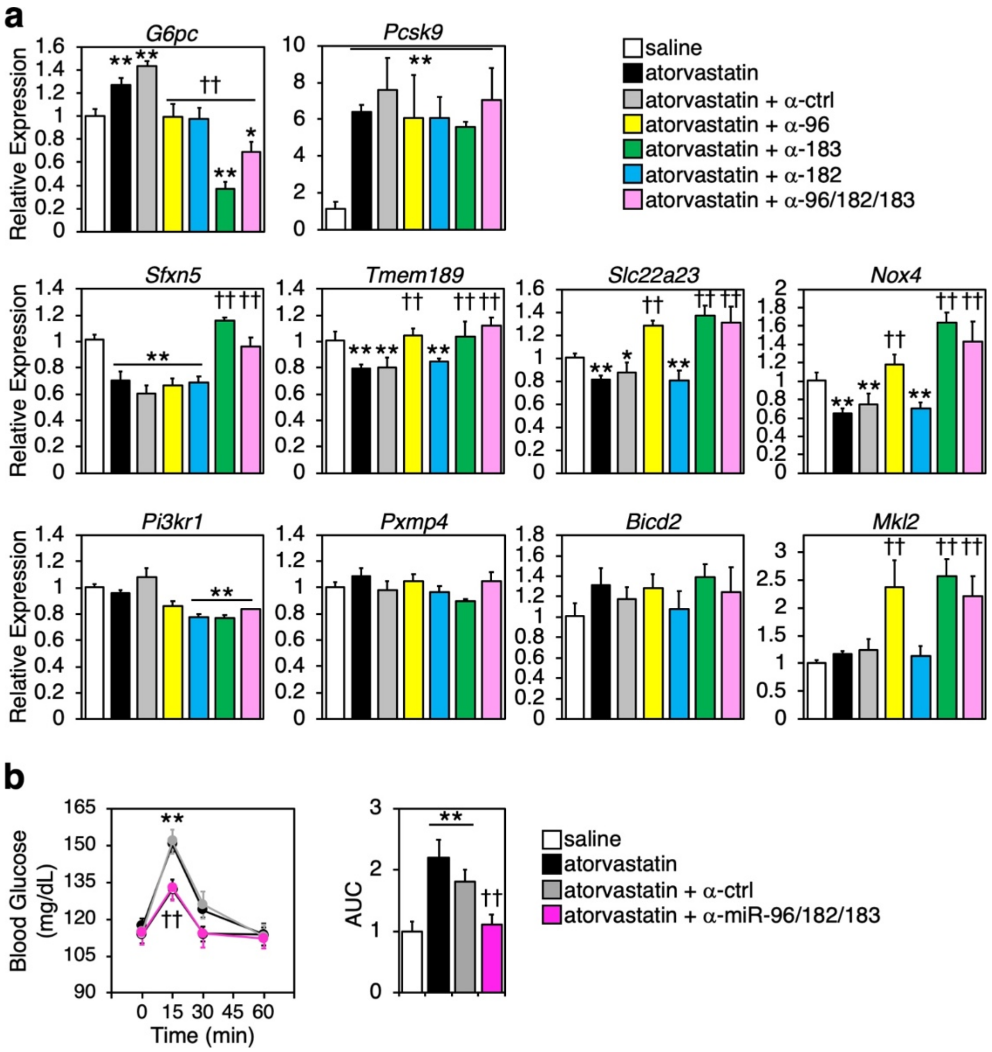


**Supplemental Figure 5. Therapeutic silencing of the miR-183/96/182 cluster abrogates statin-induced hepatic gluconeogenesis. a,** Treatment with anti-miR oligonucleotides abrogates atorvastatin-induced increased expression of hepatic *G6pc* mRNA, consistent with the reduced blood glucose in response to pyruvate challenge as shown in Fig. 4f. Panel also shows liver mRNA expression of additional miR-183/96/182 predicted targets (identified in Figs. 1e and 2c) in the same livers. **b,** A second cohort of mice (n=7/group) were gavaged with saline or atorvastatin, and then injected with anti-miRs, as described for Fig. 4d–f. Pyruvate challenge was performed as in Fig. 4f, and confirmed that therapeutic silencing of the miR cluster abolished atorvastatin-induced gluconeogenesis. Data are mean ± SE. $^{*}p<0.05$, $^{**}p<0.01$, compared to saline. $^{\dagger}p<0.05$, $^{\dagger\dagger}p<0.01$, compared to atorvastatin or [atorvastatin + anti-ctrl].
